# JOINT ACTION AWARENESS IN MACAQUES DURING A MULTIPLAYER VIDEO-GAME CONTEXT

**DOI:** 10.64898/2026.10.19.759127

**Authors:** V. Papagni, E. Quarta, S. Grasso, A. Rinaldi, A. Battaglia-Mayer

## Abstract

In behavioral neuroscience, research in nonhuman primates often relies on video-game–like paradigms, including studies of social interaction and joint action. In social studies, establishing whether animals monitor their partner and are aware of their reciprocal interactions in these abstract laboratory settings is essential for interpreting the processes underlying cooperative performance. Here, we examined spontaneous head-orienting behavior in two macaques performing a cooperative visuomotor task under highly controlled conditions. The animals sat side by side with their heads unrestrained and used isometric joysticks to control their respective cursors displayed on a shared screen. Depending on trial instructions, they performed either individual actions (SOLO action, SA) or joint actions (JA) to obtain a reward. We quantified spontaneous head rotations toward the partner as an ethologically relevant behavioral indicator of social monitoring. Partner-directed head turns occurred more frequently during JA than during SA trials. Crucially, their rate increased with the number of errors made by the partner during JA trials. Together, these findings indicate that macaques monitor their partner during abstract, computer-mediated cooperative interactions and that this monitoring is driven by the partner’s performance. Such spontaneous, context-specific orienting toward the partner supports the interpretation that macaques are aware of their dyadic engagement, even in video-game-like settings. Our results therefore highlight the suitability of controlled visuomotor paradigms for investigating the cognitive-motor processes underlying joint action in nonhuman primates.

## INTRODUCTION

Since the pioneering work of Evarts and later Mountcastle (Evarts, 1968; Mountcastle et al., 1975), macaques have served as a fundamental animal model in behavioral neuroscience. Their sensorimotor systems share remarkable anatomical and functional similarities with those of humans, making them uniquely suited for investigating the neural mechanisms underlying perception, movement control and action selection. Over the years, research in non-human primates has expanded beyond basic sensorimotor physiology to encompass higher-order cognitive processes, including decision-making, action representation and executive functions. This broader perspective has enabled the study of increasingly abstract visuomotor behaviors, such as maze solving, spatial navigation, rule-based categorization and complex sequential tasks.

Addressing these scientific questions typically requires highly structured and often sophisticated behavioral paradigms. Such hyper-controlled environments where monkeys are tested are essential not only for technical reasons, ensuring stable recordings, reproducible trial structure and clearly defined behavioral variables, but also to isolate the neural processes of interest from potentially confounding cognitive, sensory or motor events. However, this level of experimental control often produces contexts that are distant from the animals’ natural behavioral repertoire. A common critique therefore concerns the ecological validity of these paradigms: to what extent do neural or behavioral responses measured during joystick manipulation, button presses or interactions with computer screens reflect processes that macaques genuinely use in ethologically relevant conditions?

This issue becomes even more prominent when studying the neural bases of social interaction, whether within game-theoretical frameworks (Haroush & Williams, 2015; Moeller et al., 2023) or from a motor control perspective, where cooperation requires motor coordination between two agents. To investigate the dynamics of joint action, cooperative decision-making and the achievement of shared goals, we have developed abstract, video-game-like paradigms in which two monkeys sit side by side and control visual cursors using joysticks (Visco-Comandini et al., 2015; Ferrari-Toniolo et al., 2019; Lacal et al., 2022; Pezzulo et al., 2022; Quarta, Lacal et al., 2025; Sacheli, Grasso et al., 2026). These tasks offer the tight experimental control needed to disentangle individuals from cooperative cognitive–motor processes, but they also diverge substantially from natural macaque social behaviors.

This raises a central and legitimate question: are monkeys aware that they are interacting with a partner when performing these computer-based tasks, similar to a multiplayer videogame context? More specifically, can cooperative behavior observed in abstract visuomotor games be meaningfully interpreted as social interaction, or does it primarily reflect the sensorimotor computations imposed by task rules?

To address this question, we examined the behavior of two macaques trained to perform two versions of a visuomotor task, both performed in a dyadic setting (Visco-Comandini et al., 2015; Lacal et al., 2022). In both task versions, the two animals sat side by side with unrestrained heads, each controlling an independent cursor by applying forces to their respective isometric joysticks. Depending on the visual instruction cues, the monkeys had to guide their cursors either individually or jointly from a central position to a peripheral target to obtain a reward. The two versions differed only in the task phase in which the instruction to move together or alone was provided, a procedural difference that is not relevant for the purposes of the present study. As a proxy for the animal’s awareness of acting together, we measured their spontaneous head rotations (SHRs) toward the partner during task execution. These orienting behaviors may represent an ethologically relevant marker of awareness of being engaged in joint action context, during such videogame context, rather than simply responding to computer-controlled stimuli. SHRs are also relevant to highlight patterns of social monitoring of the partner’s actions during the laboratory-based interactive paradigm. We hypothesized that spontaneous partner-oriented head movements would occur more frequently in association with phases of joint engagement, when the correct behavior of both subjects was essential for successful task performance, than in conjunction with individual actions, and particularly following errors that could disrupt dyadic outcome. This prediction was based on the premise that errors constitute highly salient social events, as they involve the disruption of a shared goal and may therefore trigger increased monitoring of the partner. Social action monitoring supporting response optimization has been described in macaques with premotor and medial prefrontal circuits mediating these processes (Sliwa & Freiwald, 2017; Ninomiya et al., 2020; Lanzilotto et al., 2017), and with the medial frontal cortex specialized in monitoring others’ errors during social interactions. In agreement with this expectation, our results show that monkeys systematically oriented their head toward their partner during those task phases categorized as joint action engagement, which strictly require the cooperation of both subjects to reach the reward. These behavioral findings point toward the evidence of awareness of the dyadic and interactive nature of the task, suggesting that, even within an abstract laboratory setting, macaques actively incorporate partner-related information during cooperative performance. Our findings support the use of tightly controlled dyadic tasks to investigate the cognitive and motor bases of joint action, highlighting the capacity of non-human primates to extract socially relevant information even when tested in tasks that are apparently far removed from their natural behavior.

## MATERIALS AND METHODS

### Animals and experimental apparatus

Two adult male rhesus monkeys referred to here as Mk 1, Mk 2 (Macaca mulatta; body weight 8.5 and 9 Kg, respectively) were used in this study.

The two animals were placed together in a sound attenuated chamber and seated side-to-side on two primate chairs in front of a 40-inch monitor (100 Hz, 800×600 resolution, 32-bit color depth; monitor-eye distance: 150 cm; **Fig. 1A**). They were tested in visuomotor isometric tasks (see below for details), by using an isometric joystick, with their preferred arm (Mk 1, left; Mk 2, right) while the other was gently restrained. The behavior of the same pair of animals had been previously analysed (Visco-Comandini et al., 2015; Lacal et al., 2022) while performing the same tasks adopted in the present study, with the monkeys tested under head-restrained (HR) condition, following the standard experimental setup typical of behavioral neurophysiology, which allows for the acquisition of acute extracellular recordings and accurate eye-movement monitoring. Critically, the main analyses of the present study are instead based on sessions in which the animals performed the same behavioral tasks adopted in Visco-Comandini et al., (2015) and Lacal et al., (2022), under head-free (HF) context. Briefly, each animal was trained to control the movement of a colored circular cursor (diameter: 0.6 degrees of visual angle; DVA) displayed on a black screen, by applying a hand force on an isometric joystick (ATI Industrial Automation, Apex NC) in two dimensions on horizontal plane (sampling frequency: 1 kHz). The applied force was proportionally converted into a motion of cursors on the x and y axis of the vertical plane of the monitor. The NIH-funded software package REX was used to control stimuli presentation and to collect behavioral events and force data. During experimental procedures, all efforts were made to optimize animal welfare. Animal care and housing procedures were in conformity with European (Directive 63-2020 EU) and Italian (DL.116/92 and DL. 26/2014) laws on the use of non-human primates in scientific research. The experimental procedures were authorized by the Ministry of Health (DM 162/2010-c, 163/2010-b, 96/2014-c).

**Fig. 1.**
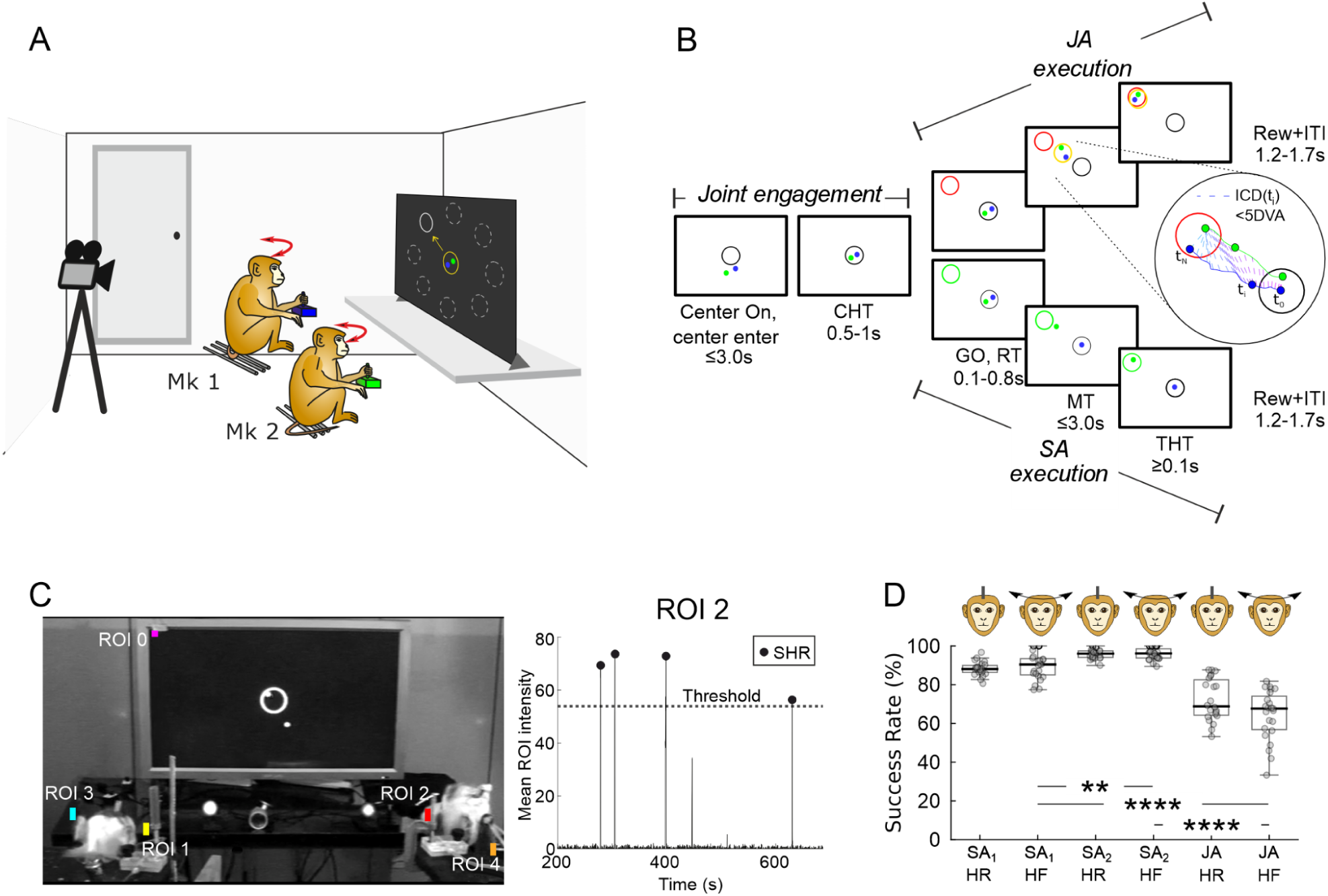
Experimental paradigm and success rates across task conditions. **(A)** Two monkeys were seated side by side in front of a common monitor, in a head-free context (HF), which allowed spontaneous head orientation toward locations capturing their attention. Each monkey guided a visual cursor (blue, Mk 1; green, Mk 2) from a central target toward a peripheral target using an isometric joystick to obtain a liquid reward, in a SOLO (SA) or a joint (JA) condition. During task performance spontaneous head rotations (SHRs) were detected via video analysis. **(B)** Description of the timeline of one of the two versions of the task (no-PI, PI version not shown; see Methods). Upon onset of the central target (Center On), both monkeys were required to move their cursors inside the central target and keep it there for a variable time (0.5 to 1s, CHT). The time from Center On to the end of the CHT is here referred to as the “Joint Engagement” phase. The GO signal cued by the appearance of a peripheral target indicated the location to be achieved by the cursor, and the action condition (SA, blue or green target or JA, red target). We referred to the epochs after the GO signal as “SA-” or “JA-execution” phase. During JA execution, monkeys had to reciprocally coordinate their movements in order to bring a common object (a yellow circle) towards the peripheral target. Inset: cursors’ trajectories of both monkeys during a typical JA trial, with dashed lines illustrating the time-resolved inter-cursor distance (ICD), to be kept below 5 DVA. **(C)** Detection of SHRs. In the video frame (left) the placement of Regions of Interest (ROIs) are indicated: ROI 0, on small light stimulus indicating trial initiation, used to temporally align the video recordings with task events; ROI 1 and ROI 2, detecting partner-directed SHRs for Mk 1 and Mk 2, respectively; ROI 3 and ROI 4 detecting Mk 1’s and Mk 2’s SHRs directed toward an alternative control location (Door). The right panel displays the mean intensity extracted from ROI 2 over time, during a representative temporal window. Following signal filtering, intensity peaks exceeding the detection threshold (dashed line) were classified as SHRs (black dots). **(D)** Comparison of success rates (SR) across action conditions (SA, JA) and across different experimental contexts (HR, HF). Statistical significance was determined using the Kruskal-Wallis test with Bonferroni post-hoc test. For visual clarity, only the pairwise comparisons between HR and HF contexts and those between action conditions tested under HF context are indicated (*, p < 0.05; **, p < 0.01; ****, p < 0.0001; no asterisk: p > 0.05).

### Behavioral task

Monkeys were required to perform an isometric center-out task in which they guided their own cursor (blue for Mk 1, green for Mk 2) from the central position to the peripheral target, by applying a dynamic hand force on their own joystick, under two conditions requiring two different action types: Solo Action (SA) and Joint Action (JA), **Fig. 1B**. Only successful trials were rewarded with liquid. All trials began with the presentation of an outlined white circle (diameter: 4 DVA) on the center of the screen (Center On). Each animal had to bring its cursor inside it and hold it there for a variable control time (Center Holding Time, CHT; 500-1000 ms). Importantly, the entrance of both cursors controlled by the two animals inside the central target and the fulfillment of the CHT were a prerequisite to continue the trial and receive the cues about the action type to perform in order to receive the reward. We termed this initial phase, including the time allowed to enter the central target (max 3 s), as “Joint engagement” (*Jeng*), since it required a mutual engagement of both subjects before they were instructed which type of action to perform. Following the Jeng phase, trials then progressed to the execution phases of either solo action (SAexec) or joint action (JAexec). The execution phase typically comprises distinct epochs (e.g., reaction time, movement time, target holding time), however, for the purposes of the present study, these epochs have been merged together, and collectively referred to as *SAexec* and *JAexec* (**Fig. 1B**). In the SA condition, the animal performed the task individually, by moving its own cursor alone toward the peripheral target in order to get reward, independently from the partner’s behavior. In the JA condition, both monkeys had to move their cursors towards the peripheral target jointly, by coordinating their hand forces in intensity direction and speed, in order to move a common visual object (represented by a yellow circle) from the center to the peripheral location (**Fig. 1B**). To instruct the monkeys about the action types to be executed we adopted two different task versions, as described in detail in previous work (Lacal et al., 2022): a no-Pre-Instructed task (noPI) and a Pre-Instructed task (PI) task. In the noPI task, the information about the action type and location of peripheral target (force direction) were provided simultaneously (**Fig. 1B**); while in the PI task the two information were provided separately and sequentially: the action condition was first pre-instructed during a dedicated variable interval, after which the directional cue was presented (not shown; see Lacal et al., 2022 for details).

Specifically, in the noPI task, at the end of the CHT, a peripheral target (outlined circle, 4 DVA in diameter) was presented at one of eight possible positions signaling the GO cue (**Fig. 1B**). Its color instructed the animals about the type of action required to obtain a liquid reward. Therefore, in the noPI task, force direction and condition were cued simultaneously. Following the appearance of the peripheral target (GO signal) the animals were required to move their own cursor (individually or together) within a given RT (100-800 ms limits) towards the new location to get the reward. Details about inter-individual coordination constraints are described below.

In the PI task, the CHT was followed by a color change of the central target that pre-instructed the monkeys about the upcoming action during the pre-instruction time (PIT, 1000-1500 ms): blue for SA_1_, green for SA_2_, and blue-green for JA. During the PIT, the animals were required to keep their cursors within the central target. When a white peripheral target (GO signal) appeared in one of eight possible positions, monkeys had to move their cursor either individually or jointly toward the target within the allowed RT (100-800 ms), according to the instruction cue. In SA trials, only the instructed monkey moved to the target to obtain a reward after holding the cursor on the target for at least 100 ms (THT), while the partner maintained its cursor at the center to obtain its own reward; the two behaviors were independent and had no effect on each other. In JA trials, both monkeys had to move their cursors toward the peripheral target together by coordinating their forces, as in the noPI condition (see below for coordination constraints).

In both PI and noPI tasks, different peripheral locations were presented in an intermingled and pseudo-randomized fashion. Each trial was followed by an inter-trial interval (ITI) of variable random duration (1200-1700 ms) to prevent anticipatory responses.

Notably, the two task versions were presented in separate sessions, under head-unrestrained conditions. A total of 2877 noPI trials and 3402 PI trials were collected.

### Inter-individual coordination constraints

In both versions of the task, JA condition required monkeys to move their cursors towards the peripheral target in a coordinated fashion to gain both the same amount of liquid reward, after holding the cursors in the final position for at least 100 ms. For the entire duration of their cursors’ motion, monkeys were constrained to maintain a maximum inter-cursor distance (ICDmax) of 5 DVA, which coincided with the size of the yellow circle encompassing the two cursors (**Fig. 1B**). The instantaneous position of the center of the yellow circle coincided with the mean value of the x and y coordinates of the two cursors. Exceeding the ICDmax resulted in an “ICD error” which led to trial abortion and none of the two agents obtained a reward.

### Detection of spontaneous head rotations

To record spontaneous head rotations (SHRs) during task performance, a video camera (25 fps, 480p) was positioned behind the two interacting monkeys (**Fig. 1A**). A total of ∼5 hours of recordings (6279 trials) was acquired across 5 days of behavioral testing in the tasks described above. For the purposes of SHR quantification and statistical analysis, the entire duration of the videos was segmented into non-overlapping ∼15-minute blocks. This interval was chosen because it matched the duration of the shortest recording, ensuring comparable analyses across videos. In this context, SHRs directed toward the partner were used as a behavioral proxy of the animal’s awareness of co-acting with its partner during this multiplayer video game-like task.

All video recordings were analyzed using custom scripts implemented in Python and MATLAB. First, original RGB images were converted to single channel grayscale images to isolate luminance information and reduce computational load (**Fig. 1C**). Then, for each time frame, the Mean Pixel Intensity (MPI) was extracted from specific Regions of Interest (ROIs, see below).

To ensure precise temporal synchronization between video-derived behavioral data and the behavioral data collected during the task, through the REX acquisition system, a light stimulus marking trial onset was detected by a photocell. The signal was extracted from the video using a region of interest (ROI 0) positioned over the stimulus in the top-left corner of the screen (**Fig. 1C**).

To detect SHRs during the task performance from the video recordings, we implemented a custom script to extract automatically the time-resolved MPI from the four ROIs (ROI 1 to ROI 4). These ROIs were positioned bilaterally near each monkey’s head and were specifically placed to detect SHRs towards the monkey’s respective partner, as shown in **Fig. 1C**. An example of this automatic detection is shown in **Movie 1**, where the MPI time series extracted from the ROIs are overlaid on the video. It is worth noting that the door of the recording chamber (located behind the monkeys on their left side) represented a potential source of distraction (e.g., auditory stimuli, coming from outside) unrelated to the behavioral task, which could therefore elicit SHRs directed toward that location. Hence, as a control, two additional ROIs (ROI 3, ROI 4) were positioned to detect SHRs toward the door of the recording room (**Fig. 1C**). Since our hypothesis specifically concerns the exchange of social cues triggered by salient events during SA or JA, the main analyses focus on the partner-directed SHRs obtained from ROI 1 and ROI 2, with the SHRs towards the door obtained from ROIs 3 and 4 used as controls.

For Mk 2, since the rotation toward its partner was in the same side compared to the rotation toward the door (leftward) we implemented an additional exclusion method to accurately disentangle SHRs directed specifically toward the partner from those directed toward the door: any MPI peak detected in the primary partner-oriented ROI 2 was disregarded if it co-occurred within a 200 ms window of a peak detected in the control ROI 4. This 200 ms temporal constraint was established by averaging the minimum inter-peak latencies derived from the signal’s first derivative. This criterion was essential to differentiate genuine social-relevant rotations from head orientations unrelated to social interaction. Automatic MPI peak detection was implemented using a dynamic thresholding technique, which calculates a video-specific threshold for each ROI to adapt to its local characteristics. First, each ROI time series was normalized by subtracting the mean MPI value of a 20-second baseline window. This baseline was manually identified via visual inspection of the periods during which the monkey observed the video without head rotations. MPI Peaks were then extracted using the ’findpeaks’ function in MATLAB, with ’MinPeakDistance’ (set to 0.5s) and ’MinPeakProminence’ as key parameters. If multiple peaks occurred within this time window, only one peak was counted. The second parameter was calculated as the average MPI plus 2.5 standard deviations (SDs) calculated during the **baseline window**. A detection threshold was then set at the 70% of the largest peak detected across the time series of each ROI, for each video. This 70% threshold was empirically validated through a systematic visual inspection of video recordings, verifying that the detected peaks rigorously corresponded to actual SHR.

### Behavioral data analysis

Since behavioral strategies during visuomotor tasks can differ between head-restrained (HR) and head-free (HF) conditions, with head-free paradigms allowing more natural eye-head coordination and orienting behavior (Freedman & Sparks, 1997), we first conducted a comparative analysis of behavioral performance in our task between HF and HR testing conditions to assess whether and to what extent the two contexts affected the performance of our task. The HF condition, which allows greater freedom of movement and potential for distraction, influences task execution compared to HR condition as a classical laboratory setting which is typically employed to facilitate precise eye-movement monitoring and acquisition of neuronal activity. For this purpose, we utilized a dataset from the same subjects while these performed our task in HR.

***Success Rate***. To quantify overall behavioral performance of each animal, for each session of the two experimental settings (HR and HF) we calculated the success rate (SR), during individual action (SA_1_ and SA_2_ for Mk 1 and Mk 2, respectively) or for the dyad in the case of JA. For the SR analysis, we considered only those trials for which the engaged animal(s) completed the CHT successfully and displayed a RT. The Kruskal-Wallis test with Bonferroni post-hoc test was employed to evaluate the effect of the experimental setting on the performance.

***Error Rate***. The ER in different error types was determined for each block, as the percentage of errors relative to the total number of trials. Statistical comparisons between the two animals across the different error types were performed using a two-way ANOVA, followed by the Tukey-Kramer post-hoc test. The individual contribution to the cooperative performance was more specifically characterized by measuring the ER of each monkey during Jeng and JAexec phases, computed as the number of errors committed by one animal, normalized by the total number of trials during these epochs, within each block. We excluded in this case ICD errors, since such shared failures preclude single-subject attribution. Significant differences in ER between monkeys were assessed using the Wilcoxon rank-sum test.

***Reaction Time and Peak of Speed***. To investigate whether the different actions (SA, JA) influenced movement kinematics, a statistical analysis was conducted by two-way ANOVA test to compare the RT and the cursor peak of speed (PS) across conditions and monkeys. In each SA and JA trial the RT was defined as the time elapsing from the presentation of GO Signal to the onset of the cursor’s motion. This movement onset was defined as the first change in the cursor velocity exceeding the mean + 3 standard deviation (SD) of the signal calculated during the 50 ms before and after the presentation of the peripheral target, for at least 90 ms. The PS was estimated for each movement, as the maximum value of the cursor’s tangential velocity. Both RTs and PSs were computed exclusively on successful trials.

***SHRs to the partner***. The analysis of SHRs was focused on exploiting these movements toward the partner as a proxy of joint action awareness in our dyadic task. We categorized SHRs as being associated with a specific trial if the head rotation occurred within a predefined temporal window encompassing the entire trial duration plus the subsequent ITI, calculated trial by trial. Specifically, a SHR was associated with a specific trial, if its onset timestamp (SHR_onset_) satisfied the following condition:

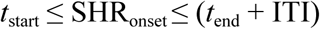

Where t_start_ and t_end_ represent the trial onset and offset, respectively. This criterion ensured that the analysis captured only those SHRs strictly associated with a specific trial’s epoch and its immediate outcome feedback.

Within the Jeng phase, to ensure that our SHRs detection approach captured only those rotations effectively reactive to a lack of engagement, we applied a temporal filter, by including in our counts only those SHRs occurring at least 600 ms after the Center On signal. This threshold was determined based on the average latency to enter the central target during successful trials (∼300 ms) and an estimated RT (∼300 ms) to initiate a head movement following error detection (failure to enter the target). Rotations occurring earlier than this threshold were considered unlikely to reflect social monitoring.

We calculated the frequency of SHRs associated with errors, occurring during the Jeng and JAexec phases, as these were considered as a cooperation failure, in contrast to SAexec errors. Statistical differences in SHRs frequency across task phases were assessed through a Kruskal-Wallis test (phase: three-level factor, followed by Dunn–Šidák post-hoc tests for pairwise comparisons). Behavioral differences between the two monkeys were also assessed by using a Wilcoxon rank-sum test.

All samples were initially tested for normality using the Kolmogorov-Smirnov test. When samples exhibited a non-normal distribution, non-parametric statistical tests were applied. Specifically, the Wilcoxon rank-sum test was employed for two-sample comparisons, and the Kruskal-Wallis test was utilized for comparisons involving k>2 groups, followed by Dunn–Šidák post-hoc tests for pairwise comparisons. Subsequently, a systematic investigation using Spearman’s correlation was performed to evaluate how the frequency of partner-directed SHRs was modulated by trial outcome.

### Modeling SHR occurrence as a function of consecutive errors

To investigate whether the occurrence of repetitive errors modulated social monitoring via SHRs, we analysed the relationships between the probability of SHRs and the number of consecutive errors (1, 2, ≥3), preceding a given SHR. This analysis was performed separately for own and partner-generated errors across all trials (regardless of the type of errors). To this aim, consecutive errors were calculated as follows: an error streak was started (count = 1) the first time a monkey committed an error, and was incremented with each consecutive error made by that same animal. For this analysis we excluded ICD errors, as these cannot be unambiguously attributed to a single animal.

The error streak was treated as “own” from the perspective of the monkey committing the error and as ’partner’ from the perspective of the monkey that did not commit the error. The streak was reset following a successful trial, if the other monkey committed an error, and by the end of each video. Each error trial yielded one observation per monkey, comprising the following information: video, error source (“own” or “partner” from the point of view of each monkey), the consecutive error count, the number of SHRs directed towards the partner (SHR_partner_) and, as a control condition, the number of SHRs towards the door (SHR_door_, the second main attractor for monkey head orientations; see above).

We then plotted, for each monkey, the probability of SHR_partner_ and the probability of SHR_door_ as a function of the number of consecutive errors across all trials. This probability was defined as the total number of trials with n (1, 2, ≥3) consecutive errors that had at least one SHR divided by the total number of trials with n (1, 2, ≥3) consecutive errors, separately for own and partner-generated errors.

For statistical inference, for each animal trial-level SHR counts were modeled using generalized linear mixed-effects models (GLMMs). This approach allowed modeling the trial-level number of SHRs as the dependent variable, the accumulated consecutive error count as a fixed effect and the video as a random intercept. Models were fitted with a Poisson error distribution and a log link function, which is appropriate for count data and enables the estimation of multiplicative effects of predictors on SHRs. All models were implemented using the ’glmer’ function from the *lme4* package (Bates et al., 2015), in R (version 4.3.3). To test whether consecutive partner or own errors specifically influenced head orientation toward the partner or the door, the data of each monkey were fitted separately to the following models:

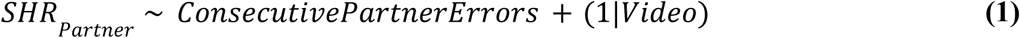

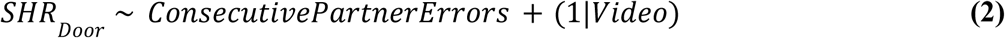

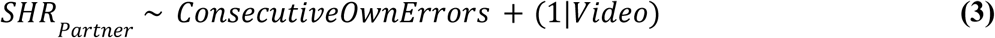

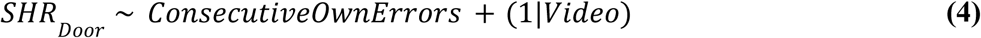

Using simulated residuals generated by the DHARMa package in R (version 0.4.7; Hartig et al., 2024), we ruled out both potential overdispersion and zero-inflation for all models (all p > 0.05). From the GLMMs, we report the incidence rate ratios (IRR) with their 95% confidence intervals (CI) and *p*-values. The IRRs represent the multiplicative effect on the expected SHR rate associated with the number of consecutive errors.

All statistical analyses were performed using custom scripts in MATLAB (MathWorks), Python or R. Unless noted otherwise, p < 0.05 was defined as the criterion for statistical significance.

## Data availability

No part of the study procedures or analyses were pre-registered prior to the research being conducted. Data and scripts for reproducing our results are available upon reasonable request.

## RESULTS

### Comparison of task performance across head-restrained and head-free conditions

Previous work has demonstrated that acting together yields a lower success rate (SR) relative to acting alone, while testing monkeys behavior under head-restrained (HR) context (Visco-Comandini et al., 2015; Ferrari-Toniolo et al. 2019; Lacal et al., 2022; Quarta, Lacal et al., 2025; Sacheli et al. 2026). While the HR condition represents the standard experimental condition in macaque behavioral neurophysiology, enabling precise neural recordings and eye-movement monitoring, it prevents the assessment of spontaneous head movements that may provide important information about the animals’ behavior, particularly when interacting in social contexts. Accordingly, our hypothesis was that by testing animal behavior in social interactions tasks under head-free (HF) condition, spontaneous head rotations could provide a behavioral proxy for the monkeys’ awareness of being engaged in a dyadic (joint-action) task. Given our interest in this new HF experimental condition, we first asked whether it produced behavioral performance different from that previously observed in the HR context.

As an initial step, we therefore compared task performance between the HR and HF contexts to assess whether limitations to head movements affected the animals’ ability to perform the task. We focused on the success rates (SR) across action types (i.e., SA and JA) performed in HR and HF experimental settings. We found that for each action type, SR remained remarkably stable across the two settings (SA_1_: 88±4% in HR, 90±7% in HF; SA_2_: 96±2% in HR, 96±3% in HF; JA: 72±11% in HR, 64±13% in HF; in all comparisons, HR vs HF, Kruskal-Wallis test followed by Bonferroni post-hoc test, p > 0.05; **Fig. 1D**). As expected, under the HF condition similarly to the HR setting, SR during JA was lower with respect to SA (SA_1_ vs JA and SA_2_ vs JA, Kruskal-Wallis test, followed by Bonferroni post-hoc test, p < 0.0001; **Fig.1D**). Moreover, when comparing the SR of the two monkeys during their own SA, we found that Mk 1 had lower SR compared to Mk 2, similarly to the case of HR context (**Fig. 1D**, Bonferroni post-hoc test, p < 0.01).

Our findings thus indicate that monkeys performed the task equally well under both HR and HF conditions, with comparable success rates across settings and with the typical decrease in SR when acting jointly compared to when acting alone. Inter-individual differences between the two animals in SR during solo actions were also maintained. In light of these results, and given the focus of the present study on head movements, subsequent analyses were restricted to the HF condition.

### Subject-specific error rates and movement kinematics in head-free monkeys

Motivated by the performance differences observed between monkeys during solo actions, and based on the hypothesis that errors may play a key role in eliciting attention toward the partner, we next quantified the frequency of different error types (error rates, ER), irrespective of action condition (JA or SA), to identify which errors were most commonly committed by each animal (**Fig. 2A**).

**Fig. 2.**
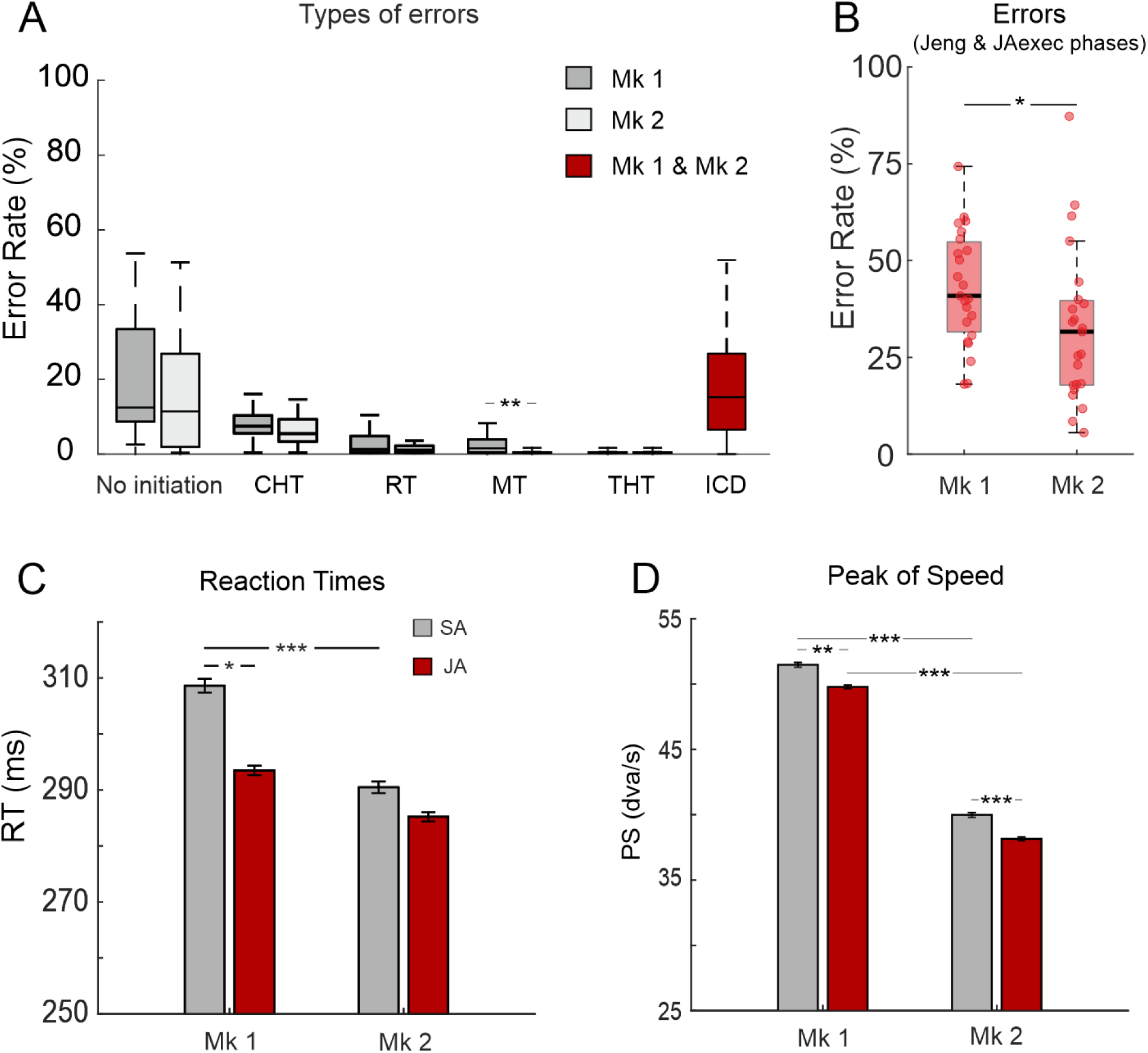
**Error rates and movement kinematics in head-free monkeys**. **(A)** Error rates for different error types: No Initiation, CHT (Center Holding Time), RT (Reaction Time), MT (Movement Time), THT (Target Holding Time), and ICD (Inter-Cursor Distance). Error percentages were calculated relative to the total number of trials for each monkey separately (Mk 1, dark gray; Mk 2, light gray), except for ICD errors, which reflect shared coordination failures that cannot be attributed to either monkey individually (two-way ANOVA, Tukey-Kramer post hoc test; no asterisk, p > 0.05; **p < 0.01). **(B)** Individual error rates calculated both during Joint engagement (Jeng) & JA execution (JAexec) phases (Wilcoxon rank-sum test, *, p < 0.05). **(C)** Reaction Times (RT, mean (SE) in ms) during SA (gray) and JA (red) trials (two-way ANOVA, Tukey-Kramer test, *p < 0.05; ***p < 0.001). **(D)** Peak Speed (PS, mean (SE) in dva/s) during SA and JA trials (two-way ANOVA, Tukey-Kramer test **, p < 0.01; ***, p < 0.001).

ER analysis revealed that the two subjects shared a similar distribution of errors across all error types (two-way ANOVA, Monkey × Error Type interaction: F_(4,230)_=42,4, p < 0.0001; Tukey-Kramer post-hoc test: p > 0.50) except for the one related to the MT error, which occurred more frequently in Mk 1 compared to Mk 2 (3±3% vs. 1±1.6%, respectively; Tukey-Kramer post-hoc test: p =0.001).

In both animals, the most frequent errors corresponded to trial initiation failures (here termed as ’No initiation’), occurring when the monkeys did not initiate the trial after the initial instruction cue., i.e when they did not enter the central target within the requested 3 s time window. Furthermore, a specific error of the JA condition, consisted in the so-called ’ICD errors’, caused by an inter-cursor distance exceeding the ICDmax during task execution (see Methods), which contributed heavily in affecting the outcome of the cooperative performance (16.6±14%), as also reported in previous studies performed in head-restrained monkeys (Lacal et al., 2022).

Given the social nature of the task, we focused on the potential differences in performance between co-agents during cooperation. We thus characterized each monkey’s contribution to cooperative performance by analyzing their individual error rates (ER) during the Jeng and JAexec epochs. We computed the overall ER during these epochs for each monkey, excluding ICD errors, as these represent shared coordination failures for which responsibility cannot be unambiguously attributed to a single animal. This analysis revealed that Mk 1 committed more such errors compared to Mk 2 (Mk 1: 45±17%, Mk 2: 34±22%, Wilcoxon rank sum test, p < 0.05, **Fig. 2B**).

Beyond reduced overall performance when cooperating as quantified by the SR (or ER), our previous studies testing HR animals in similar task conditions to the present study (Visco-Comandini et al., 2015; Ferrari-Toniolo et al., 2019; Lacal et al., 2022; Quarta, Lacal et al., 2025; Sacheli, Grasso et al., 2026) provided evidence for proactive adjustments in movement kinematics when acting together respect to individual actions. In the present study, we asked whether such adjustments would persist in the two animals, also in the new HF setting. We therefore focused our analysis on the reaction times (RT) and on the peak movement speed (PS) as representative of the planning and execution phase, respectively.

When tested under the HF setting, monkeys exhibited kinematic adaptations (**Fig. 2 C-D**) to solve the JA task, similar to those previously observed under HR conditions (Visco-Comandini et al., 2015; Lacal et al., 2022; Quarta, Lacal et al., 2025; Sacheli, Grasso et al., 2026).

We first observed that MK1 was significantly slower than MK2 when performing the task under the SA condition (two-way ANOVA, main effect Monkey: F_(1,3719)_ =758.22, p < 0.0001), and that this animal shortened its RT when acting jointly (SA_1_: 309 ± 98 ms; JA: 293 ± 82 ms; Tukey-Kramer post hoc test, p < 0.001), achieving RTs similar to those of its partner. MK2, which was already faster during SA, did not significantly change its RT between SA and JA (SA2: 290 ± 67 ms; JA: 285 ± 65 ms; Tukey-Kramer post hoc test, p > 0.05). Statistical comparisons between the animals’ RTs confirmed no significant difference during JA (Tukey-Kramer post hoc test, p > 0.05; **Fig. 2C**).

We examined the speed of force application, by analysing the peaks of speed (PS). We observed an overall significant effect of action condition (two-way ANOVA, main effect Condition: F_(1, 3719)_ = 17.55, p < 0.0001): both animals significantly reduced their movement speed during JA relative to SA (Mk 1: 51 ± 14 vs 50 ± 14 dva/s; Mk 2: 40 ± 11 vs 38 ± 11 dva/s; Tukey-Kramer post-hoc p < 0.01). During successful cooperation we found a significant inter-individual PS difference (two-way ANOVA, main effect Monkey: F_(1,3719)_ = 758.22, p < 0.0001), with Mk 1 being significantly faster than Mk 2 during JA (Mk 1: SA1 vs Mk 2: SA2, Tukey-Kramer post-hoc p < 0.001; **Fig. 2D**).

These results provide evidence for subject-specific contribution towards cooperative performance, showing that in our monkey pair, one subject (Mk 1) committed more errors than the other. In addition, this animal showed more adaptive behavior during JA, shortening its RTs so that movement initiation became more closely aligned with that of the partner, thereby fulfilling the interpersonal coordination requirements of the task.

### Quantification of spontaneous head rotations during task performance

We initially investigated potential relationships between overall outcome of the task and social monitoring, to determine in particular whether trial success or failure elicited different levels of attention toward the partner. To this aim the spontaneous head rotations (SHRs) directed toward the partner were quantified, in relation to trial outcome (successful vs error trials). We found that the vast majority (93%) of SHRs toward the partner were associated with error trials, i.e., occurring between trial onset and the end of the ITI (see Methods, **Fig. 3A**). This was statistically confirmed at individual-level in both animals (Mk 1: success 0.1±0.3 SHRs vs errors 4±3.6 SHRs; Mk 2: success 0.1±1.3 SHRs vs. errors 10±8, Wilcoxon rank-sum test, both p < 0.001; **Fig. 3B**). Furthermore, a direct comparison between the two monkeys revealed a significant difference in the frequency of SHRs performed during error trials, with Mk 2 showing a significantly higher rate of partner-directed rotations compared to Mk 1 (Mk 1: 4±3.6 SHRs vs. Mk 2: 10±8 SHRs, Wilcoxon rank-sum test, p < 0.01, **Fig. 3B**). Importantly, the higher frequency of SHRs in Mk 2 relative to Mk 1 cannot be attributed to a greater task disengagement (e.g., increased distractibility leading to head orientation toward the partner), as this animal was more accurate than Mk 1, committing fewer errors during JA (**Fig. 2B**).

**Fig. 3.**
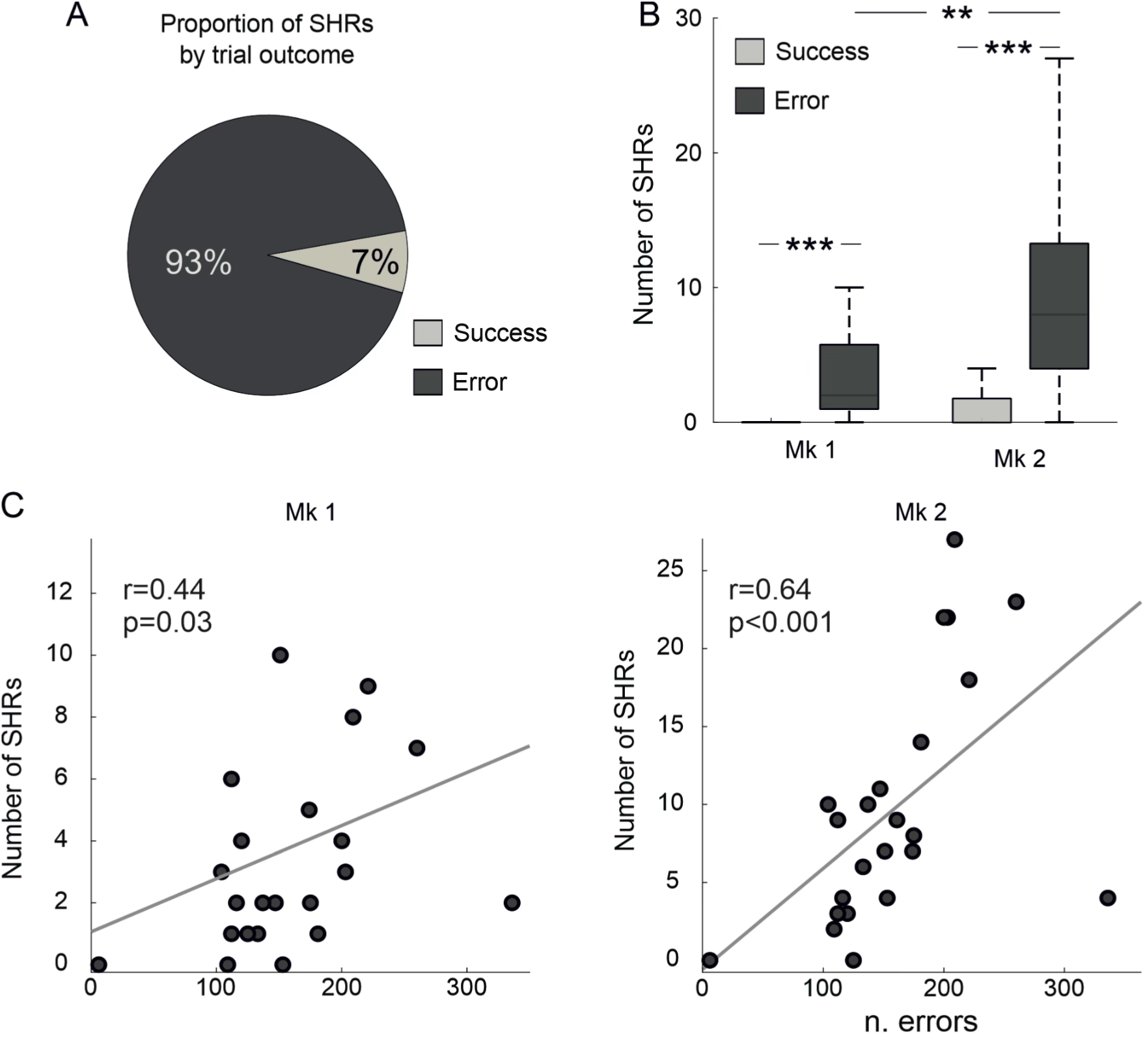
**Relationship between partner-oriented SHRs and error trials**. **(A)** Proportion of SHRs toward the partner associated with different trial outcomes (success and error), pooled across monkeys. **(B)** Comparison of SHR proportions between monkeys, separately for success and error trials (Wilcoxon rank-sum test, **p < 0.01; ***p < 0.001). **(C)** Spearman’s correlation between the number of error trials and the number of partner-oriented SHRs following errors, independently on action condition. Each dot represents data for a single 15-min block.

Since SHRs occurred mostly within unsuccessful trials, we investigated whether the frequency of SHRs toward the partner was indeed related to the error rate. In both monkeys, we found that the number of error trials significantly and positively correlated with the frequency of SHRs (Mk 1, Spearman’s r=0.44, p=0.03; Mk 2, r=0.64, p < 0.001, **Fig. 3C**).

Together, these findings suggest that when macaques act in a dyadic context, the frequency of their spontaneous head orientation towards the partner is highly influenced by the number of behavioral errors. Rather, this difference may be related to the greater number of errors committed by Mk 2’s partner, a relationship examined in detail below.

So far, we have not distinguished the action context (SA or JA) in which the errors were committed by the two animals. Our paradigm in fact distinguishes trials based on whether the cursor must be moved individually (SA) or jointly (JA). However, despite this distinction, in both cases trials share an initial phase preceding the GO signal, beginning with the appearance of the central target, during which both monkeys had to move their cursors inside it, and keep it for a variable CHT, as a prerequisite for trial continuation. Then, at the end of CHT, the visual cue appeared to instruct the animals about the action type to be executed (SA_1_, SA_2_, or JA; see Methods; **Fig. 1B**). Therefore, the initial phase of the trial, regardless of whether it would subsequently unfold as a SA or JA condition, required the engagement of both subjects to allow task continuation. We defined this period as a “joint engagement” (Jeng) phase, which has to be fulfilled jointly for successful task performance. Crucially, failures during this phase likely constitute highly salient social events, as they affected the payoff of both monkeys . Thus, we hypothesized that Jeng errors may elicit social monitoring that could be captured via SHRs. Consistent with this view, both monkeys exhibited the highest frequency of SHRs during trials with errors occurring in the Jeng phase compared to those with errors committed within SA and JA execution epochs (Kruskal-Wallis test followed by Dunn–Šidák post-hoc test, p < 0.001; **Fig. 4A**). No significant differences emerged when comparing SHRs between the SA and JA execution epochs (Dunn–Šidák post-hoc test, p > 0.05, **Fig. 4A**).

**Fig. 4.**
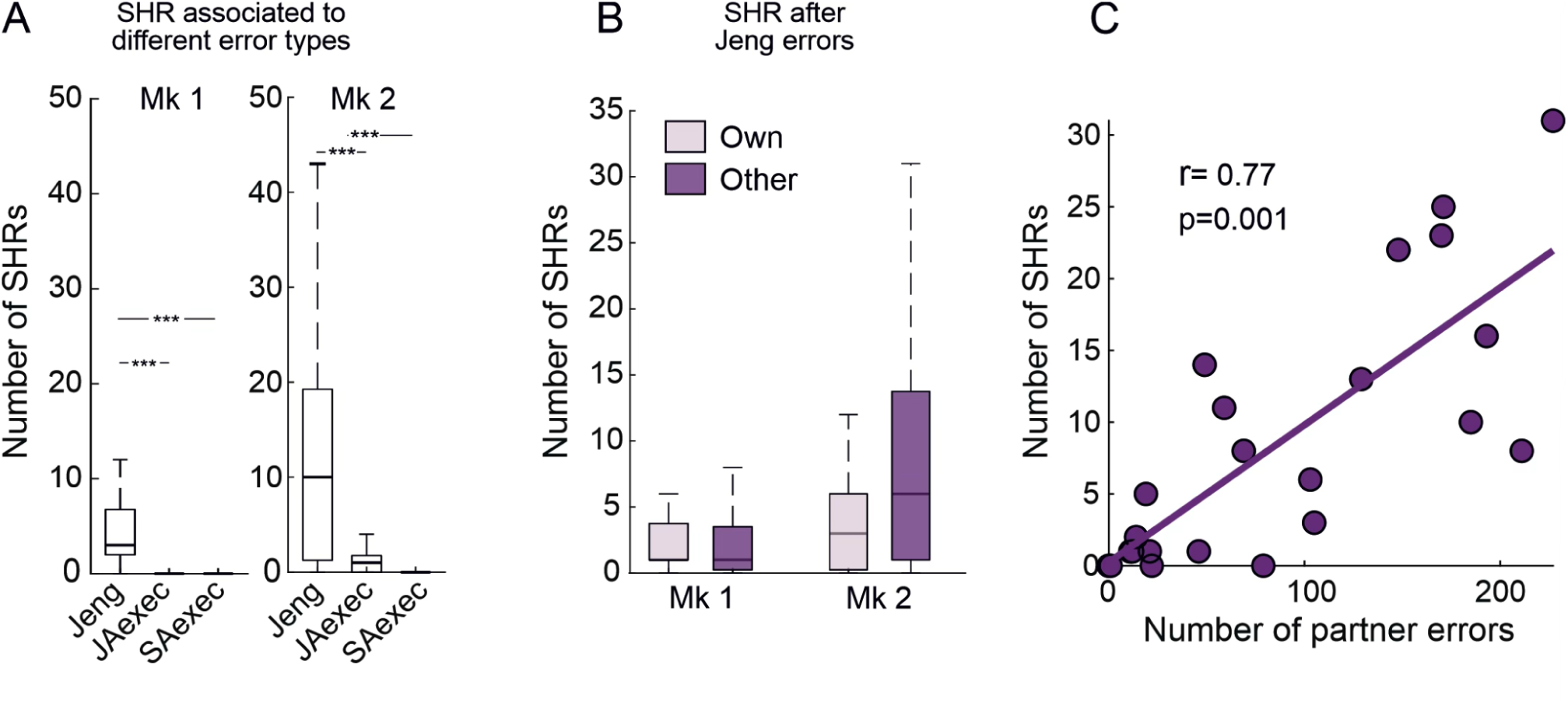
SHRs across different error types. (**A**) Number of SHRs calculated per block during error trials, across the different task phases: Joint engagement (Jeng), JA execution (JAexec) and SA Execution (SAexec). Kruskal-Wallis test, followed by Dunn–Šidák post-hoc test (***, p < 0.001). (**B**) Number of SHRs computed for self-generated (“Own”) and partner’s errors (“Other”) during the Jeng phase. No statistical differences were detected (Wilcoxon rank-sum test; no asterisk, p > 0.05), but with significantly greater variability for partner-generated errors observed in Mk 2 (Brown–Forsythe test, p < 0.05). (**C**) Spearman’s correlation in Mk 2 between the number of the partner’s Jeng errors and the frequency of SHRs toward the partner.

Given that SHRs are predominantly associated with Jeng errors, we investigated whether head orientations toward the partner depended on the error source (“own” vs “partner”). We found no significant differences in mean SHR frequency between error sources in either monkey (Wilcoxon rank-sum test, p > 0.05; **Fig. 4B**), but a significant greater variability in Mk 2’s SHR in concomitance of the partner errors, compared to all other monkey × error-source combinations (Mk1: “own” and “partner”; Mk2: “own”; Brown–Forsythe test, p < 0.001, post-hoc Tukey-Kramer test). In light of this higher variability, we next examined whether SHR occurrences covaried with the number of partner-generated errors. This analysis was restricted to data of Mk2, as the low variability of SHRs in the other monkey would limit variance and preclude reliable estimation of correlations. A strong positive correlation between SHR occurrence and the number of errors committed by the other agent (Spearman’s r = 0.77, p = 0.001; **Fig. 4C**) emerged, indicating that the probability of head orientations toward the partner increases with the partner’s level of disengagement from the joint action.

Finally, we analysed the probability of SHRs toward the partner as a function of the number of consecutive errors. We found a higher probability of partner-oriented head rotations with increasing numbers of consecutive partner errors, an effect that was not observed for head movements toward the control location (door; see Methods, **Fig. 5A**). Indeed, this trend was confirmed by generalized linear mixed-effects models: in Mk 1, trials with three or more consecutive partner errors were associated with a significant increase in SHR rate (IRR = 4.67, 95% CI [1.56–14.03], p = 0.006), and a similar effect was observed in Mk2 (IRR = 2.10, 95% CI [1.39–3.16], p < 0.001). In sharp contrast, consecutive errors did not predict head orientations toward the control area (door) in either monkey (all p > 0.05), indicating that this effect is specific to partner-directed orienting rather than reflecting a general increase in head movements.

**Fig. 5.**
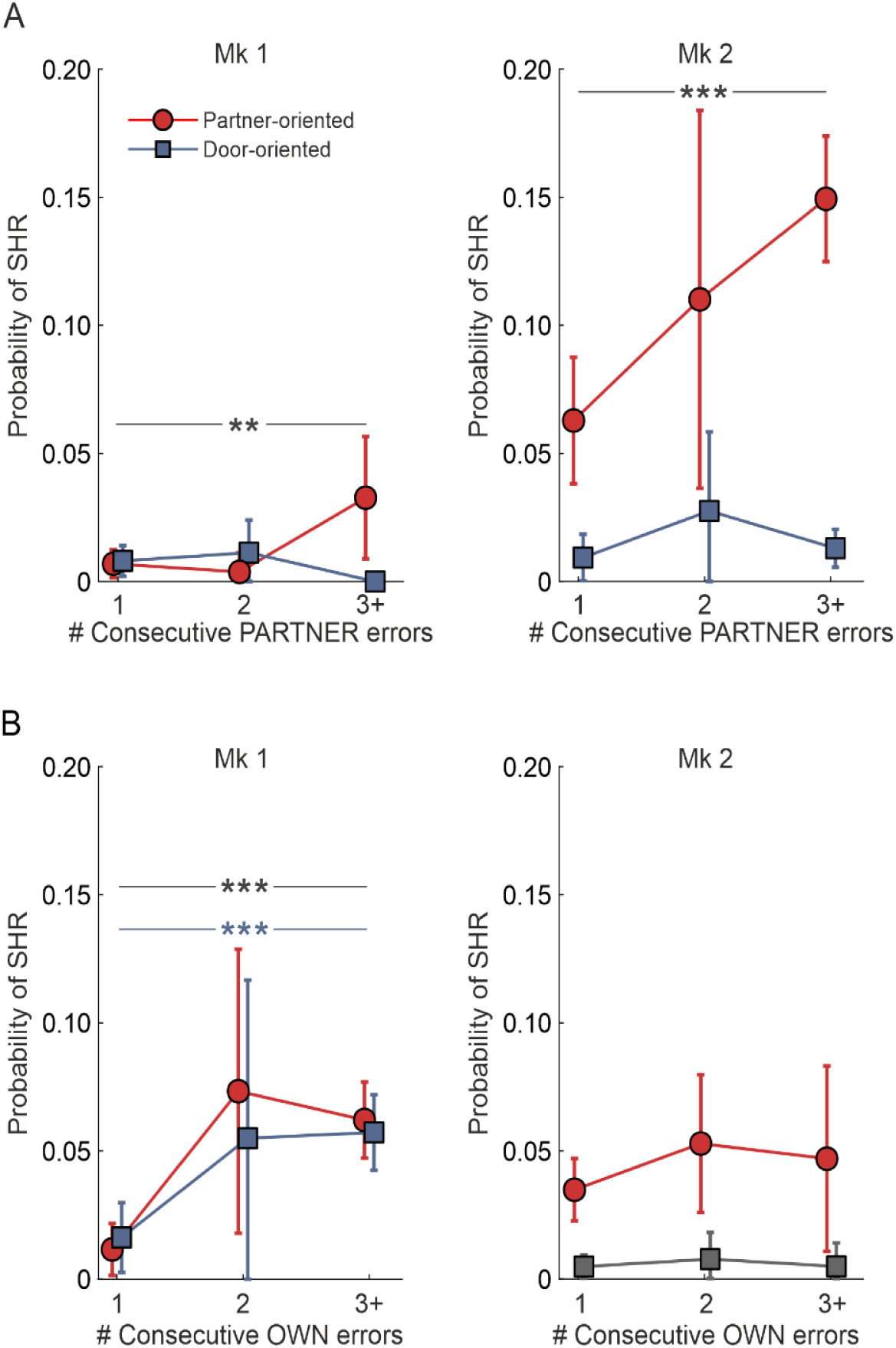
**Probability of partner-directed SHR following consecutive partner errors**. **(A)** Probability of SHR toward the partner (red circles) and toward the control location (door; blue squares) as a function of the number of consecutive partner errors (1, 2, ≥3) preceding head orientation for Mk 1 and Mk 2 (left and right, respectively). **(B)** Probability of SHRs, toward the partner (red) and the door (blue) as a function of the number of preceding consecutive self-generated (“own”) errors for each monkey for Mk 1 and Mk 2 (left and right, respectively). Data points and error bars represent empirical sample means and 95% confidence intervals calculated across individual trial observations. Horizontal bars and brackets indicate statistically significant effects and contrasts derived from the GLMMs (* p < 0.05, ** p < 0.01, *** p < 0.001).

It is worth noting that this pattern was specific to errors committed by the partner, as analyses of own errors did not show similar trends, being qualitatively different and not consistent across monkeys. In fact, in Mk1, head orientations following its own errors showed a comparable probability of SHRs occurring toward both locations. Whereas, in Mk2, SHR frequency did not increase with the number of consecutive own errors (**Fig. 5B**). Consequently, these data do not support a systematic relationship between own errors and partner-directed orienting, precluding clear conclusions about their association.

Together, these findings indicate that repeated partner errors selectively drive partner-directed orienting, in contrast to self-generated errors that do not exert a comparable influence on social monitoring behavior.

## DISCUSSION

The fundamental advances in behavioral neurophysiology using animal models have relied on highly controlled laboratory paradigms to investigate the neural bases of cognition and action. A key feature of these approaches is the use of head restraint setup (HR), enabling stable intracortical recordings and precise control of eye movements. While ensuring technical reliability and control over confounding variables, these paradigms often impose abstract task structures that do not fully capture the natural repertoire of primate behavior (Tremblay et al., 2023), particularly in the social domain. To contrast this limitation, more interactive and dynamic frameworks have recently been introduced (Berger et al., 2020; Tremblay et al., 2023; Franch et al., 2024; Lanzarini et al., 2025).

In the context of motor control during social interactions, we developed a multiplayer, video game-like paradigm, to investigate the neural bases of joint action (JA) in monkeys (Visco-Comandini et al., 2015; Lacal et al., 2022; Ferrari-Toniolo et al., 2019; Pezzulo et al., 2022; Sacheli, Grasso et al., 2026). The experimental set-up was designed to balance the needs for strict experimental control with that of developing a task requiring an online motor coordination during real-time interaction with a partner. From a neurophysiological perspective, this experimental framework has enabled us to characterize the multiscale neural coding of JA in the macaque premotor cortex, at both the single-neuron (Ferrari-Toniolo et al., 2019; Pezzulo et al., 2022; Sacheli, Grasso et al., 2026) and local field potential (LFP) levels (Sacheli, Grasso et al., 2026), while also allowing a direct comparison of behavioral and neural processes across primate species (Satta et al 2017; Fanghella et al 2025; Sacheli, Grasso et al., 2026). In particular, Sacheli, Grasso and collaborators, using the same JA paradigm adopted in the present study, combined fMRI recordings in humans with LFP recordings in macaques, revealing an evolutionarily conserved premotor mechanism for interpersonal motor coordination (Sacheli, Grasso et al., 2026). However, the use of such an abstract paradigm in non-human primates raises a fundamental question: are macaques indeed aware of the social nature despite its non-ethological context?

The primary objective of the present study was to identify behavioral signatures of awareness of acting together in such a non-naturalistic paradigm. To this end, we tested animals under head-free conditions, enabling the assessment of spontaneous, ethologically relevant behaviors, such as head rotations toward the partner, as potential indicators of social monitoring and awareness of shared engagement during task execution. Allowing spontaneous head movements may be particularly relevant in the context of dyadic interactions, as the interplay between action execution and partner-viewing has been shown to drive learning to cooperate (Franch et al., 2024; Parodi et al., 2025).

Additionally, comparing animal behavior within the HF and HR settings using an otherwise identical experimental paradigm provided a secondary opportunity to evaluate how experimental constraints influence task performance. This comparison may also provide information about animal welfare considerations during data collection, by offering insight into how different levels of physical constraint affect animal behavior and, indirectly, may indicate difference in animal distress.

### Comparison between HF and HR conditions

Our HF model showed no significant differences in success rates (SR) compared to the HR condition, when executing both solo and joint actions, indicating comparable task performance across settings. Similarly to previous HR studies (Visco-Comandini et al., 2015; Ferrari-Toniolo et al., 2019; Lacal et al., 2022; Quarta, Lacal et al., 2025; Sacheli, Grasso et al., 2026), also when the head was free to move, we observed a similar drop in performance, during JA compared to solo trials. Furthermore, although a direct quantitative comparison was not the primary aim of this study, the movement kinematics observed in the HF condition exhibited patterns of behavioral modulation across action types (SA and JA) similar to those previously reported in HR macaques. Establishing this consistency between HR and HF contexts was essential to ensure that both paradigms rely on comparable task engagement and coordination demands. Notably, the similarity in performance across conditions suggests that the head-restrained setup, at least within the constraints of the present task, is not associated with overt behavioral signs of distress sufficient to compromise animal behavior, particularly given the extensive training and progressive habituation of the animals.

### Spontaneous Head Rotations during JA performance

The analysis of spontaneous head rotations toward the partner revealed that macaques correctly interpret abstract on-screen stimuli, i.e. cursor motion, as representing the partner’s actions. We observed that partner-oriented SHRs occurred predominantly in association with error trials, consistent with the role of error monitoring as a fundamental cognitive process, to detect deviations from intended goals (Fu et al., 2023). Moreover, action monitoring not only supports adaptive behavior during online motor control of one’s own actions (Archambault et al., 2009, 2011), but also plays a key role in social contexts, when the action goals are shared with others (Musco et al., 2023). For example, neurophysiological evidence from human musical duets has shown that performers monitor not only the outcomes of their own actions but also those of their partner and of their combined actions (Loehr et al., 2013). Furthermore, studies in humans have shown that similar neural mechanisms underlie the processing of both self-generated errors and errors committed by others (van Schie et al., 2004). At the single-neuron level, studies in monkeys have further revealed distinct neuronal populations in the medial frontal cortex selectively signaling self-generated versus partner-generated errors (Yoshida et al., 2012). These mechanisms may allow individuals acting in social environments to distinguish between their own errors and those committed by others, thereby supporting accurate error attribution.

More specifically, the majority of SHRs observed in our study were temporally associated with the errors occurring during the initial phase of joint action, when both monkeys were required to bring their cursors into the central target to ensure trial continuation. This temporal distribution suggests that partner-directed monitoring is particularly prominent during early coordination phases, when the shared motor plan is at risk of failure, mutual engagement is essential. Interestingly, similar behaviors have been documented in humans engaged in collective actions. For example, in musical ensembles, bidirectional gaze between performers has been observed at the beginning of musical pieces, potentially facilitating mutual engagement and temporal coordination (Bishop, 2018). Moreover, visual monitoring of preparatory gestures has been shown to facilitate synchronization at movement onset (Bishop & Goebl, 2018). Together, these results support the interpretation that macaques engage in partner-oriented error monitoring, particularly during early phases of joint action, even when interaction is mediated by abstract visual signals.

Our results showed that in both animals, the probability of SHRs toward the partner increased as a function of the number of consecutive errors committed by the partner. This phenomenon can be regarded as a behavioral index of evidence accumulation (Steinhauser & Yeung, 2012; Pereira et al., 2021), where observing repetitive consecutive errors increases animal confidence that its partner is responsible for the trial failure. In line with this view, neural systems involved in error processing have been shown to integrate incoming signals reflecting accumulated evidence, ultimately supporting the emergence of awareness (Wessel et al., 2011).

A critical element supporting the social specificity of this orienting behavior is provided by the comparison with the SHRs oriented toward a non-socially relevant location, i.e. the door used as a control condition, indicating that the observed head movements reflect a selective coagent-oriented, monitoring process, rather than a generalized response to errors. Moreover, the relationship between SHR probability and consecutive errors did not consistently emerge for self-generated errors, further supporting the interpretation that these head turns are specifically tuned to socially relevant information.

In addition, it is worth noting that SHRs occurred relatively rarely overall. Given that executing an SHR requires diverting visual attention away from the screen, these movements can be considered a costly behavior that, while potentially enabling information accumulation (Drugowitsch et al., 2012), directly disrupts the monkey’s ongoing eye-cursor coupling. As such, orienting toward the partner is selectively deployed when the potential benefits of acquiring information about the partner outweigh the costs of interrupting ongoing visuomotor control. This interpretation is consistent with the observed increase in SHR probability as a function of consecutive partner errors. While further tests are needed to fully address the purpose of monkeys SHRs, we speculate that this behavior might extend beyond a signal of joint action awareness. Rather, these movements might embody an active communicative signal for JA (Pezzulo et al., 2026), possibly aimed at soliciting the partner to re-engage in the task and/or to avoid errors. Indeed, in natural settings, macaques might employ mutual gaze to establish and dissolve joint action engagements (Pedruzzi et al., 2025). The present findings demonstrate that such spontaneous orienting behaviors persist even in highly controlled laboratory environments, providing novel insight into their ethological relevance for joint action.

By probing joint action awareness through social monitoring, the present findings complement previous reports demonstrating that macaques exhibit understanding of self-agency, self-awareness (Couchman, 2011; Chang et al., 2017), as well as partner-dependent decision-making (Ballesta & Duhamel, 2015). Overall, by showing that macaques can map abstract on-screen interactions onto their partner’s physical motor behavior, our findings not only provide further support for the existence of a “we-representation” in cooperating monkeys (Lacal et al., 2022), but also suggest that macaques exhibit awareness of acting together, even in non-ethological settings.

## ACKNOWLEDGMENTS

This study was supported by the Ministry of University and Research, Italy, grant nr. B53D23014490006 to ABM, grant nr. B53D23030110001 to ABM, Young Researcher 2024 grant nr. B83C24003140006 to EQ and by Sapienza University of Rome, Italy, grant nr. RP125199C37E290D to ABM, grant nr. B87G22001200001 to EQ. We thank Roberto Caminiti for his useful comments and suggestions during the preparation of the manuscript. We thank Denise Tomassetti for help with animal husbandry.

## AUTHOR CONTRIBUTIONS

**ABM:** Designed the experiment and collected the data. **ABM, VP, and EQ:** Designed the data analysis. **VP and EQ:** Performed the behavioral data analyses, with contributions from AR and SG. **EQ:** Conceptualized and implemented the video analysis pipeline for SHR detection and synchronization with behavioral data. Performed statistical modeling using generalized linear mixed models (GLMMs). **VP, EQ, and ABM:** Wrote the first draft of the manuscript. **All authors:** Revised and approved the final version of the manuscript.

## DECLARATION OF INTERESTS

The authors have no competing interests or other interests that might be perceived to influence the interpretation of the article.

## Notes

### Competing Interest Statement

The authors have declared no competing interest.

